# Glyphosate and its formulations Roundup Bioflow and RangerPro alter bacterial and fungal community composition in the rat caecum microbiome

**DOI:** 10.1101/2021.11.19.468976

**Authors:** Robin Mesnage, Simona Panzacchi, Emma Bourne, Charles A Mein, Melissa J Perry, Jianzhong Hu, Jia Chen, Daniele Mandrioli, Fiorella Belpoggi, Michael N Antoniou

**Affiliations:** Gene Expression and Therapy Group, King’s College London, Faculty of Life Sciences & Medicine, Department of Medical and Molecular Genetics, Guy’s Hospital, London, SE1 9RT, UK; Ramazzini Institute, Via Saliceto, 3, 40010 Bentivoglio, Bologna, Italy; Genome Centre, Barts and the London School of Medicine and Dentistry, Blizard Institute, London E1 2AT, United Kingdom; Department of Environmental and Occupational Health, Milken Institute School of Public Health, George Washington University, Washington, DC.; Department of Genetics and Genomic Sciences, Icahn School of Medicine at Mount Sinai, New York, NY, USA; Department of Environmental Medicine and Public Heath, Icahn School of Medicine at Mount Sinai, New York, NY, USA

## Abstract

The potential health consequences of glyphosate-induced gut microbiome alterations have become a matter of intense debate. As part of a multifaceted study investigating toxicity, carcinogenicity and multigenerational effects of glyphosate and its commercial herbicide formulations, we assessed changes in bacterial and fungal populations in the caecum microbiota of rats exposed prenatally until adulthood (13 weeks after weaning) to three doses of glyphosate (0.5, 5, 50 mg/kg body weight/day), or to the formulated herbicide products Roundup Bioflow and RangerPro at the same glyphosate-equivalent doses. Caecum bacterial microbiota were evaluated by 16S rRNA sequencing whilst the fungal population was determined by ITS2 amplicon sequencing. Results showed that both fungal and bacterial diversity were affected by the Roundup formulations in a dose-dependent manner, whilst glyphosate alone significantly altered only bacterial diversity. At taxa level, a reduction in Bacteroidota abundance, marked by alterations in the levels of *Alloprevotella, Prevotella* and *Prevotellaceae UCG-003*, was concomitant to increased levels of Firmicutes (e.g., *Romboutsia, Dubosiella, Eubacterium brachy group or Christensenellaceae)* and Actinobacteria (e.g., *Enterorhabdus, Adlercreutzia*, or *Asaccharobacter*). *Treponema* and *Mycoplasma* also had their levels reduced by the pesticide treatments. Analysis of fungal composition indicated that the abundance of the rat gut commensal Ascomycota *Kazachstania* was reduced while the abundance of *Gibberella, Penicillium, Claviceps, Cornuvesica, Candida, Trichoderma* and *Sarocladium* were increased by exposure to the Roundup formulations, but not to glyphosate. Altogether, our data suggest that glyphosate and its Roundup RangerPro and Bioflow caused profound changes in caecum microbiome composition by affecting the fitness of major commensals, which in turn reduced competition and allowed opportunistic fungi to grow in the gut, in particular in animals exposed to the herbicide formulations. This further indicates that changes in gut microbiome composition might influence the long-term toxicity, carcinogenicity and multigenerational effects of glyphosate-based herbicides.

## Introduction

Glyphosate is a broad-spectrum herbicide active ingredient and the most used pesticide worldwide. Glyphosate-based herbicides are used to control weeds in agricultural fields and urban environments, but also to desiccate crops shortly before harvest. The use of glyphosate-based herbicides such as Roundup has increased exponentially since their introduction at the end of the 1970s due to the wide-scale adoption of glyphosate tolerant genetically modified crops, especially in North and South America (Benbrook 2016). It is estimated that about 700,000 tons of glyphosate are used worldwide annually (Maggi et al. 2019). Although the use of glyphosate is expected to further increase by 2025 (Maggi et al. 2019), its application is reaching a plateau in some countries due to the spread of glyphosate-resistant weed species (Heap 2014). In addition, controversies surrounding the toxicity of glyphosate has led to bans or restrictions of glyphosate usage (Robinson et al. 2020).

While the carcinogenic effects of glyphosate have been demonstrated in laboratory animal studies, the overall human health effects of glyphosate are still not fully known. Numerous studies have reported toxicity from glyphosate in different organs in a large range of animal species (Mesnage et al. 2015). Glyphosate toxicity in mammals can be explained by its ability to induce oxidative stress through the alteration of mitochondrial function (Mesnage et al. 2015). Glyphosate is known to affect mitochondrial respiratory chain functions (Olorunsogo 1990; Pereira et al. 2018; Wise et al. 1992), and this can be linked to an increased production of reactive oxygen species (Bailey et al. 2018; Gomes and Juneau 2016). Oxidative stress induced by glyphosate and its commercial herbicide formulations can provide an explanation for observed genotoxic outcomes both in vitro (Benbrook 2019) and in vivo (Portier 2020) model system. Other toxicological properties of glyphosate includes possible estrogen receptor activation (Mesnage et al. 2017) and epigenetic (DNA methylation) changes leading to alteration in gene expression (Duforestel et al. 2019; Woźniak et al. 2020).

More recently, concerns have also been raised about the ability of glyphosate to have adverse effects through its interactions with the community of bacteria residing in the digestive tract, known as the gut microbiome. The toxicity of glyphosate in plants is caused by an inhibition of 5-enolpyruvylshikimate-3-phosphate synthase (EPSPS) of the shikimate pathway, causing a shortage in aromatic amino acid biosynthesis (Boocock and Coggins 1983). Although this mechanism of action does not exist in mammalian cells, microorganisms dwelling on the surface and within the gastrointestinal tract of mammalian bodies can be sensitive to glyphosate inhibition when they are equipped with the shikimate pathway (Mesnage and Antoniou 2020; Mesnage et al. 2021c). The number of studies investigating the effects of glyphosate on microbial communities is growing (Tsiaoussis et al. 2019). Our recent study using a combination of metabolomics and shotgun metagenomics has made clear that glyphosate can inhibit the shikimate pathway in the gut microbiome, causing an accumulation of metabolites upstream of EPSPS (Mesnage et al. 2021c). However, whether glyphosate effects are different between males and females, if they can be detected from exposure to low regulatory permitted levels, different across the different sections of the gastrointestinal tract (e.g., ileum, caecum, colon), or even if early-life exposure can cause more damage remains known.

Although there have been a number of reports on the effects on glyphosate and glyphosate-based herbicides on gut bacterial microbiota, no studies to date have investigated to see if there are alterations in the gut fungal population, sometimes referred to as the gut ‘mycobiome’. The role of the mycobiome is more elusive mainly because they have been under-studied compared to bacteria (Huseyin et al. 2017). This is despite growing evidence linking the presence of different categories of fungi in the human gut to observable diseases and health symptoms, such as multiple sclerosis and other neurological diseases, no study has investigated if glyphosate can affect these fungal populations (Huseyin et al. 2017).

Furthermore, commercial glyphosate herbicides do not only contain glyphosate in water, but also other ingredients called co-formulants. These co-formulants are normally listed on packaging as “inert” since they proposed to not have any direct herbicidal action. The major co-formulants used in the manufacture of Roundup herbicides are surfactants, which are used to allow glyphosate penetration through the waxy surface of plant leaves (Mesnage et al. 2019). Surfactants generally form 5-15% of the concentrated products, which has to be diluted to different concentrations depending on the intended use. For instance, Roundup (MON 2139) is used as a 2 percent solution (7.2 g/L of glyphosate, 3.6% surfactant MON 0818) on most perennial weeds. The main surfactants for several decades used in Roundup formulations have been polyoxyethylene tallow amine (POEA). More recently, different types of surfactants have been introduced, which have now very different toxicity profiles (Mesnage et al. 2019).

In a previous investigation, we described the changes in faecal microbiome composition in Sprague-Dawley rats orally exposed to glyphosate and Roundup Bioflow (MON 52276) at a glyphosate equivalent dose corresponding to the US chronic Reference Dose (cRfD) of 1.75 mg/kg bw/day starting from prenatal life until adulthood (13 weeks) (Mao et al. 2018). We found that faecal microbiota profiling revealed changes in overall bacterial composition in pups at postnatal day 31, corresponding to pre-pubertal age in humans (Mao et al. 2018). The study presented here constitutes a follow-up to this earlier investigation and forms part of the multifaceted *Global Glyphosate Study*, which was launched with the aim of providing the most comprehensive evaluation of glyphosate-based herbicides covering long-term toxicity, carcinogenicity and multi-generational effects. Glyphosate and two commercial formulations, namely the EU representative formulation Roundup BioFlow (MON 52276) and the US formulation RangerPro (EPA 524-517), were again administered to Sprague-Dawley rats starting mid-gestation via drinking water at 0.5, 5, and 50 mg/kg bw/day, which ranged from the EU acceptable daily intake (ADI) to the EU no observed adverse effect level (NOAEL). RangerPro contains POEA surfactants (Mesnage et al. 2021a), while Roundup Bioflow does not contain POEA (Mesnage et al. 2021b), however the complete co-formulant profile remains unknown. Reported here are the effects of these three compounds on the caecum microbiota.

## Materials and Methods

### Animal experimental setup

The study was conducted following the rules established by Italian law regulating the use and humane treatment of animals for scientific purposes (Legislative Decree No. 26, 2014. Implementation of the directive n. 2010/63/EU on the protection of animals used for scientific purposes. - G.U. General Series, n. 61 of March 4^th^, 2014). Before commencement, the protocol was examined by the Internal Ethical Committee of Ramazzini Institute for approval. The protocol of the experiment was also approved and formally authorized by the ad hoc commission of the Italian Ministry of Health (ministerial approval n. 945/2018-PR). All animal study procedures were performed at the Cesare Maltoni Cancer Research Centre/Ramazzini Institute (CMCRC/RI) (Bentivoglio, Italy).

The CMCRC/RI animal breeding facility was the supplier of the Sprague-Dawley (SD) rats. Female SD rats used for breeding were placed individually in polycarbonate cages (42×26×18cm; Tecniplast Buguggiate, Varese, Italy) with a single unrelated male until evidence of copulation was observed. After mating, matched females were housed separately during gestation and delivery. Newborns were housed with their mothers until weaning. Weaned offspring were housed, by sex and treatment group, not more than 3 per cage. Cages were identified by a card indicating: study protocol code, experimental and pedigree numbers, dosage group. The cages were placed on racks, inside a single room prepared for the experiment at 22 °C ± 3 °C temperature and 50 ± 20% relative humidity. Daily checks on temperature and humidity were performed. The light was artificial and a light/dark cycle of 12 h was maintained.

During the experiment animals had *ad libitum* access to an organic pellet feed “Corticella bio” supplied by Laboratorio Dottori Piccioni Srl (Piccioni Laboratory, Milan, Italy). In addition, the animals drank fresh municipal tap water from glass bottles *ad libitum*. Both feed and water were periodically analyzed to identify possible chemical or microbiological contaminants or other impurities; these analyses are included in the documentation of the experiment. The pelleted feed was tested for possible glyphosate contamination in compliance with Commission Regulation (EU) No 293/ 2013 [maximum residue levels (MRLs) < 1 mg/kg]. Tap drinking water was tested for possible glyphosate contamination in compliance with Directive 2008/105/EC, D.Lgs. 152/2006, Directive2006/118/EC (active substances in pesticides, including their relevant metabolites, degradation and reaction products < 0.1 μg/l).

Virgin female SD rats (18 weeks of age) were cohabited with an outbred male of the same age and strain. Each day, the females were examined for presence of sperm. Gestational day (GD) 0 was set on detection of sperm in vaginal smears. The day on which parturition was completed was designated as lactating day (LD) 0 for the dam and PND 0 for the offspring. After weaning (~PND 24-28), the offspring, identified by ear punch according to the Jackson Laboratory system (Enclosure 2), were housed in the same treatment group as their dams in order to have no more than one male and one female per group.

The treatment included 10 experimental groups: one untreated control group; 3 groups treated with glyphosate 3 groups treated with Roundup Bioflow® and 3 groups treated with Ranger Pro®. Test substances were diluted in drinking water at the same glyphosate equivalent concentrations such that the animals’ daily intake corresponded to 0.5 mg/kg bw day (ADI Europe); 5 mg/kg bw day and 50 mg/kg bw day (NOAEL Europe) of glyphosate.

Every animal in the experiment was checked 3 times per day on weekdays, and twice on Saturday and Sunday/public holidays. Water and food consumption, and body weight were recorded periodically. At the end of the treatment period, at 17 weeks of age (13 weeks after weaning), each surviving animal was anesthetized by inhalation of a mixture CO_2_/O_2_ (70% and 30% respectively) and sacrificed by drawing blood via *vena cava*. During necropsy, the caecum content from each animal was collected in a polypropylene tubes and stored at −80°C until use.

### DNA extraction

DNA was extracted from 0.25g of caecum content material using the PowerFecal protocol (Qiagen, Hilden, Germany) as per the manufacturer’s instructions. Extracted DNA was quantified using a Qubit instrument (Thermo Fisher Scientific, MA, USA). DNA concentrations were standardised to 5ng/μl using an automated protocol on a BiomekFX liquid handling robot (Beckman Coulter, CA, USA).

### 16S rRNA and ITS2 amplicon sequencing

PCR was performed using the Roche High-Fidelity PCR System (Roche Life Science, Welwyn Garden City, UK). A total of 5ng DNA was amplified in a reaction volume of 10μl. The primers for the amplification of the 16S V3-V4 region were: ACACTGACGACATGGTTCTACACCTACGGGNGGCWGCAG (forward) and TACGGTAGCAGAGACTTGGTCTGACTACHVGGGTATCTAATCC (reverse). The primers for ITS2 amplification were: TCTACACTCGTCGGCAGCGTCAGATGTGTATAAGAGACAGGCATCGATGAAGAACGCAGC (forward) and GTCTCGTGGGCTCGGAGATGTGTATAAGAGACAGTCCTCCGCTTATTGATATGC (reverse). The PCR reaction mixture included 1μl of 10X FastStart High Fidelity Reaction buffer, 0.1 μl of 10 μM forward or reverse primers, 1.8 μl of 25mM MgCl_2_, 0.5 μl dimethyl sulfoxide (DMSO), 0.2 μL of 10mM PCR Grade Nucleotide Mix, 0.1 μl of 5U/μl FastStart High Fidelity Enzyme Blend and 5.2 μl of nuclease-free water. This was added to 1μl of diluted samples and amplified for 35 cycles at 95°C for 30s, 55°C for 30 s, 72°C for 30s, and a final extension at 72°C for 5 minutes. The size of amplified products was verified by electrophoresis on a 2% agarose gel. We then used 1 μl of a 100 times diluted PCR product in 1xTE buffer in a second round of PCR to add TSP FLD barcodes and Illumina adaptors onto PCR products. The barcoding reaction mix included 1μl of 10X FastStart High Fidelity Reaction buffer, 1.8 μl of 25mM MgCl_2_, 0.5 μl of DMSO, 0.2 μL of 10mM PCR Grade Nucleotide Mix, 0.1 μl of 5U/μl FastStart High Fidelity Enzyme Blend and 3.4 μl of nuclease-free water. This was added to 2 μl of Fluidigm Barcode and 1μl of the 1:100 harvested PCR product. The amplification was done for 15 cycles at 95°C for 15s, 60°C for 30 s, 72°C for 60s, and a final extension at 72°C for 3 minutes. Barcode attachment was controlled using the Tapestation D1000 instrument (Agilent, CA, USA).

An equal volume of each barcoded PCR product was pooled and the final mixture diluted to 4 nM. The pooled library was loaded onto a 300bpx2 paired-end MiSeq (Illumina, CA, USA), as per the manufacturer’s instructions generating an average of 52,850 ± 10,795 reads per sample for the 16S rRNA sequencing data, and 46,883 ± 8,039 reads per sample for the ITS2 sequencing data.

### Bioinformatics analyses

The DADA2 pipeline (v 1.16) was used to quantify amplicon sequence variants (ASV) using R v4.0.0. Primers were removed using Cutadapt v2.6 with Python v3.7.1. Pseudo-pooling of samples was performed to increase the sensitivity of the analysis. The taxonomy was assigned using the native implementation of the naive Bayesian classifier method from DADA2 with the SiLVA ribosomal RNA gene database v138 for the 16S reads and the General Fasta release files from the UNITE ITS database for fungal taxonomy. Cleaned read counts, ASV taxonomies, and the metadata were then combined for an analysis with the phyloseq package v1.32.0 (McMurdie and Holmes 2013).

Alpha diversity was calculated as the number of observed species and Shannon indexes with phyloseq. Statistical significance was determined with ANOVA tests followed by Tukey post-hoc tests for multiple comparisons. Since phylum abundance was not normally distributed, statistical significance of differential phylum abundance was determined with a Kruskal-Wallis test followed by Dunn’s test of multiple comparisons. Microbiome beta diversity was compared between each sample using non-metric multi-dimensional scaling (NMDS) plots of Bray-Curtis dissimilarities, with the statistical significance of sample clustering evaluated with a permutational ANOVA (PERMANOVA) analysis on the Bray-Curtis dissimilarities with adonis from vegan v2.4-2. Sample clustering was evaluated using unsupervised PCA on centred log-ratio transformed data with microViz v0.7.1. Taxonomy was plotted using microViz v0.7.1 and ggplot2 v3.3.0. Because the gut microbiota profiles were different between males and females, we used sex as a covariate. Faecal microbiome community structure was analysed at different taxonomic levels by aggregating the content of the phyloseq object. Differences in faecal microbiome composition at the genus level were evaluated by determining multivariable association between clinical metadata and microbial meta’omic features using MaAsLin2 v0.99.12 in R with sex as a covariate. Sex-specific analyses were also performed. Data were normalised using the total scale sum and transformed with an arcsine square root transformation to account for the unevenness of read counts and stabilise the variance. The statistical significance has been corrected for multiple comparisons and presented as False Discovery Rate (FDR). Raw data are available at the NCBI sequence read archive under the accession number BioProject PRJNA758037 for the bacterial composition data and BioProject PRJNA758026 for the fungal composition data.

## Results

We measured the changes in bacterial and fungal composition in the caecum microbiota of rats exposed to three doses of glyphosate (0.5, 5, 50 mg/kg body weight/ day) or the formulated herbicide products Roundup Bioflow and RangerPro at the same glyphosate-equivalent doses. After removing bad quality reads and chimeras, we obtained 16,653± 3,947 reads per sample of bacterial 16S rRNA sequencing data, and 10,449± 2,138 reads per sample of fungal ITS2 sequencing data. Bacterial profiles from the caecum microbiome were 63% of Firmicutes (essentially *Lactobacillus, Romboutsia, Lachnospiraceae*) and 14.9% of Bacteroidota (mostly *Prevotella, Alloprevotella*), while more than 99% of the fungal species identified were from the phylum Ascomycota (*Kazachstania)*. Some Archaea represented by *Methanobrevibacter* were also detected at low abundance by the 16S rRNA sequencing platform.

Both fungal and bacterial diversity was affected by the glyphosate herbicide formulations, though in opposite directions (Figure 1). We found a statistically significant difference in average bacterial Shannon alpha diversity by treatment with both Roundup formulations (adj-p = 9.8e-06) and sex (adj-p = 6.24e-09). These pesticide treatments affected fungal alpha diversity (adj-p = 9.87e-11), but with no difference between sexes (adj-p = 0.83). Tukey post-hoc tests revealed a dose dependent decrease in bacterial Shannon alpha diversity, which was statistically significant at the highest doses for Roundup Bioflow (adj-p = 0.002) and RangerPro (adj-p = 0.004). In contrast, exposure to Roundup Bioflow caused a significant increase in fungal Shannon alpha diversity at the highest dose (adj-p = 0.00009). Exposure to RangerPro caused a remarkable dose-dependent increase in Shannon diversity, which was statistically significantly different from the lowest dose tested (adj-p = 0.03). Glyphosate alone had no effect on fungal or bacterial alpha diversity (Figure 1) suggesting that the co-formulants present in the Roundup formulations were responsible, either alone or in combination with glyphosate, for the changes observed in alpha diversity resulting from exposure to these herbicides.

**Figure 1.**
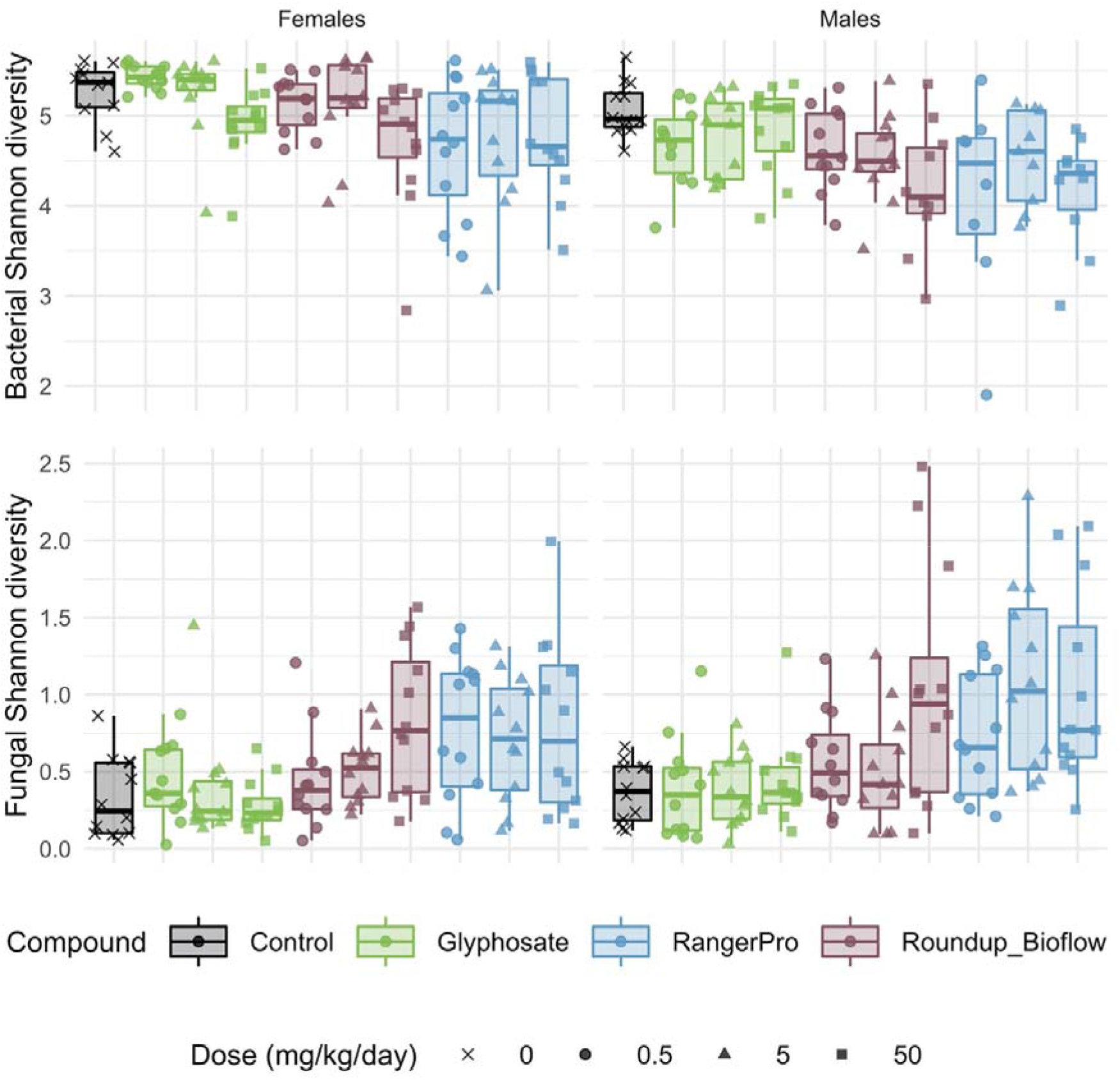
Bacterial and fungal diversity is altered in the gut microbiome of rats exposed to either glyphosate or its herbicide formulations Roundup Bioflow and RangerPro. Bacterial and fungal diversity was evaluated by 16S rRNA and ITS2 amplicon sequencing, respectively in male and female SD rats exposed via drinking water to three doses of glyphosate (0.5, 5, 50 mg/kg body weight per day), or to the formulated products Roundup Bioflow and RangerPro at the same glyphosate-equivalent doses. Exposure was initiated prenatally at mid-gestation and continued until 13 weeks post-weaning.

Beta diversity was also affected by the exposure to glyphosate herbicide formulations whilst glyphosate alone had limited effects, as demonstrated by PERMANOVA analysis revealing that both fungal and bacterial composition differ by sex (p = 0.001) and treatment groups (p = 0.001). Further analysis of sample clustering using an unsupervised principle component analysis (PCA) showed that the bacterial composition of samples from rats exposed to either Roundup Bioflow or RangerPro segregated from the control and glyphosate treatment groups (Figure 2). The PCA analysis for the fungal diversity was less clear due to the relatively low number of taxonomic groups detected, with samples clustering along the first component (Figure 2). The source of variation from the fungal microbiota sample clustering was unknown.

**Figure 2.**
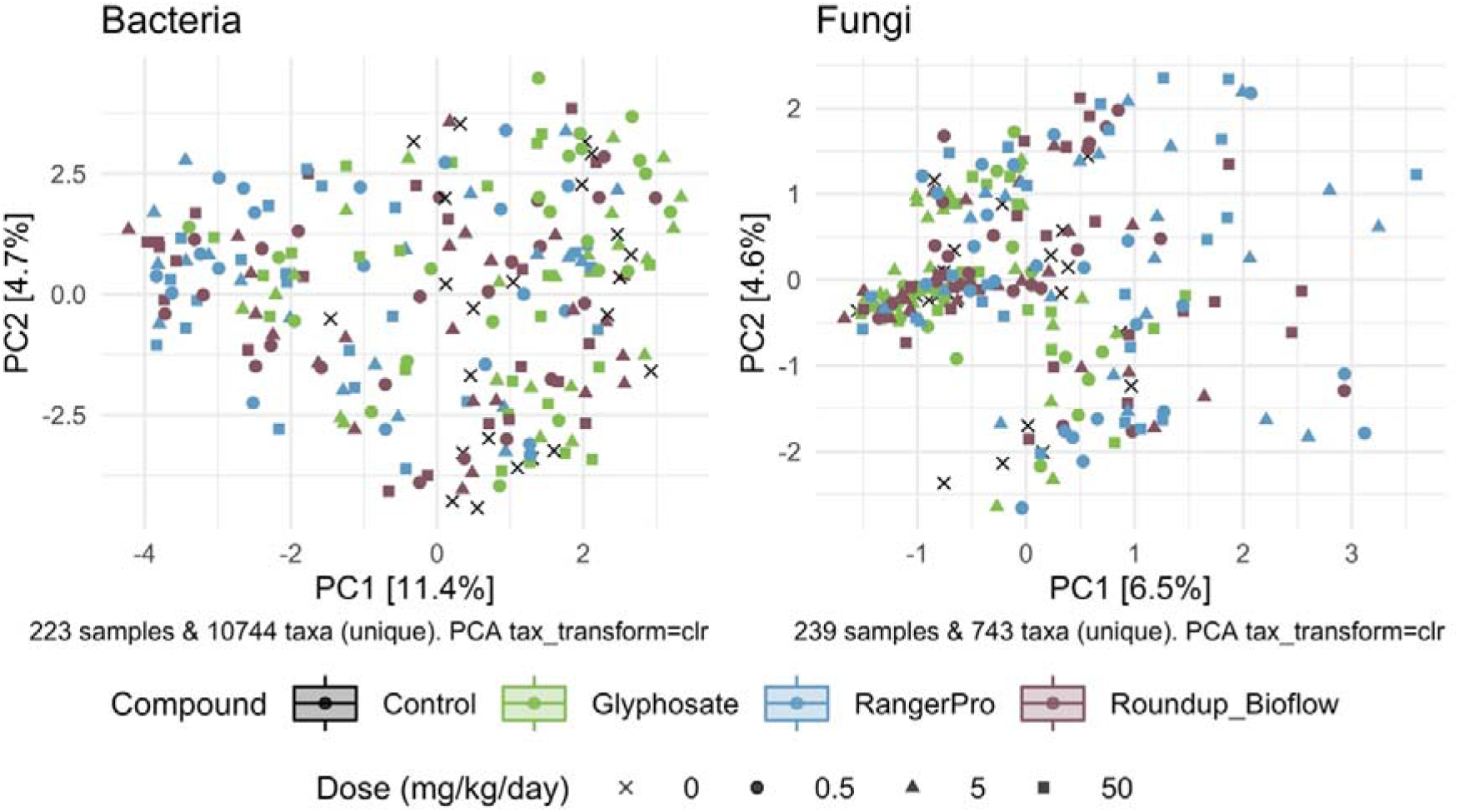
Unsupervised classification of variations in bacterial and fungal community composition by PCA. Male and female Sprague Dawley rats were exposed to three doses of glyphosate (0.5, 5, 50 mg/kg body weight per day), or to the formulated products Roundup Bioflow and RangerPro at the same glyphosate-equivalent doses starting at mid-gestation and ending at 13 weeks post-weaning. The PCA was calculated using centred log-ratio transformed abundance data from bacterial 16S rRNA and fungal ITS2 amplicon sequencing of the caecum microbiota.

We then analysed bacterial and fungal taxonomic composition to provide insight into what microorganisms were altered by exposure to glyphosate, Roundup Bioflow, or RangerPro. We found large sex differences in response to the different test substances at the phylum level for the bacterial profiles (Table 1). The most marked changes were found in females, where the abundance of Euryarchaeota was increased in a dose-dependent manner by the pesticide treatments, reaching 7.4 ± 10.2% for glyphosate at 50 mg/kg bw/day, 9.8 ± 9.1% for Roundup Bioflow at 50 mg/kg bw/day and 6.0 ± 12.0% for RangerPro at 50 mg/kg bw/day of the total assigned abundance, while Euryarchaeota only represented 0.4 ± 1.1% of the 16S assigned abundance in the control group. The same trend was visible in male animals but changes were not statistically significant. This can be due to the higher Euryarchaeota abundance in the male control animals compared to females (Table 1). In contrast, the abundance of Campilobacterota and Bacteroidota decreased in response to glyphosate treatment in a dose-dependent manner, which was again more pronounced in females than in males. Although Bacteroidota represented 24.7 ± 6.7% of the total abundance in control female animals, these species reached an average abundance level as low as 9.9 ± 7.1% for animals exposed to the highest dose of RangerPro. No changes in the levels of Basidiomycota and Ascomycota were found in any of the treatment groups (Table 1).

**Table 1.**
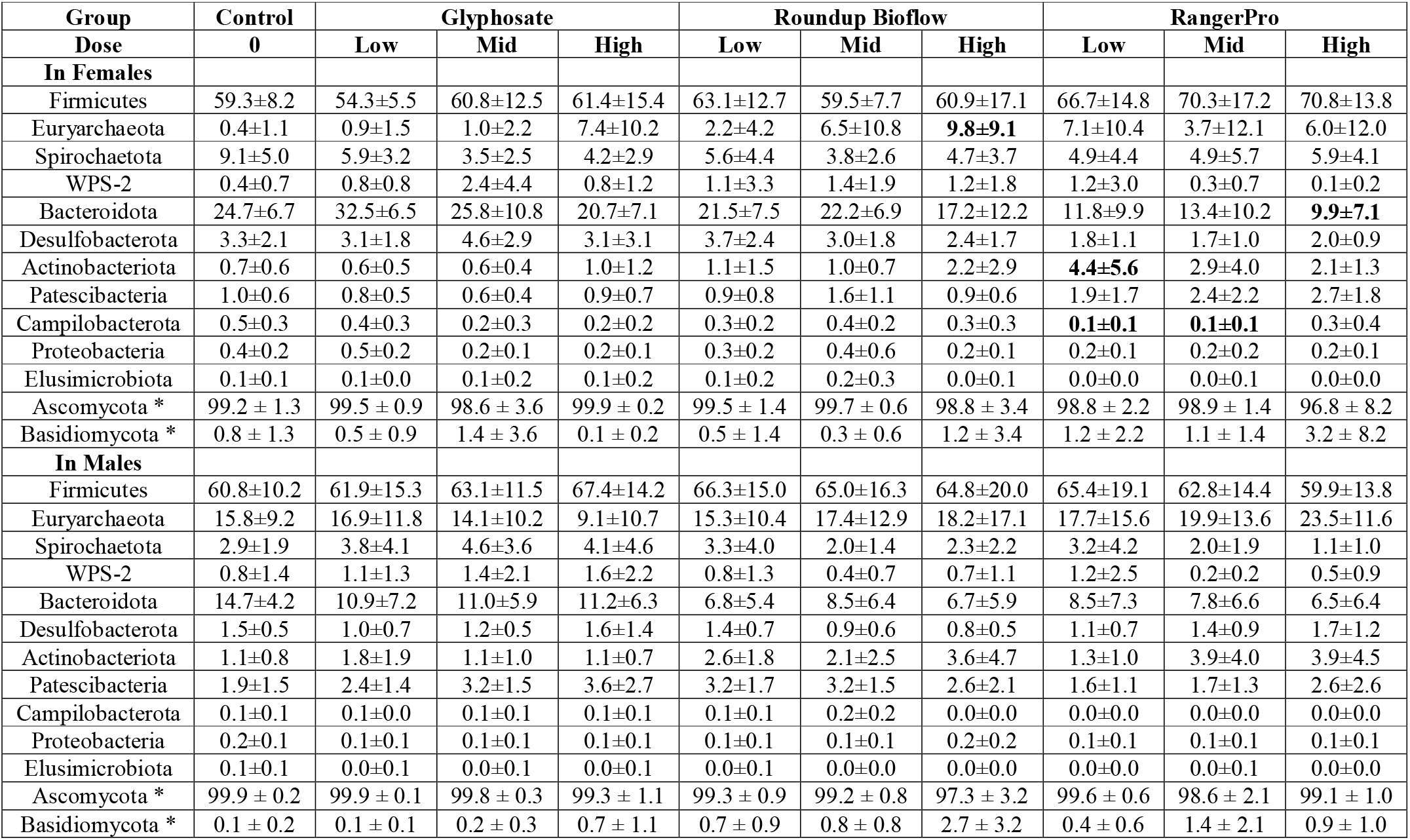
Changes in phyla abundance in the caecum microbiota of male and female rats exposed to glyphosate and the glyphosate formulated products Roundup Biolow and RangerPro. All phyla presented were those detected using the 16S rRNA platform to identify bacterial species. * The two fungal phyla detected (Ascomycota and Basidiomycota). Doses of glyphosate administered to animals: Low, 0.5mg/kg bw/day; Mid, 5mg/kg bw/day; 50mg/kg bw/day. Total abundance is presented as mean ± SD. Bold characters indicate taxonomic groups, which have their levels statistically different from the control group with a Kruskal-Wallis test with Dunn’s test of multiple comparisons.

Changes in microbial composition were also investigated at the genus level. Although DADA2 provided taxonomic assignment at species levels, a large number of sequence variants had their taxonomy unknown at the species level. Thus we decided to use genus level assignments. Given the large number of genera identified, we used a multivariate method taking into account the sex of the animals as a covariate. A large number of bacterial genera had their levels altered by glyphosate and its two herbicide formulations (Figure 3). The taxonomic composition of the most frequently found genera is presented in Figure 3A. The same genera were consistently altered by either glyphosate alone or the two Roundup formulated products (Figure 3B), although with varying degrees of statistical significance, suggesting that these disruptions originate in the ability of glyphosate to alter gut microbial metabolism. The largest of these changes reflected the overall alteration in the Bacteroidota population, marked by a reduction in the levels of *Alloprevotella, Prevotella* and *Prevotellaceae UCG-003. Treponema*, which is a genus of spiral-shaped bacteria (Spirochaetes), and *Mycoplasma*, also had their levels reduced by the glyphosate treatments. In contrast, the bacterial genera, which increased in abundance after exposure to glyphosate or its formulations were mostly Firmicutes (e.g., *Romboutsia, Dubosiella, Eubacterium brachy group or Christensenellaceae)* and Actinobacteria (e.g., *Enterorhabdus, Adlercreutzia*, or *Asaccharobacter*).

**Figure 3.**
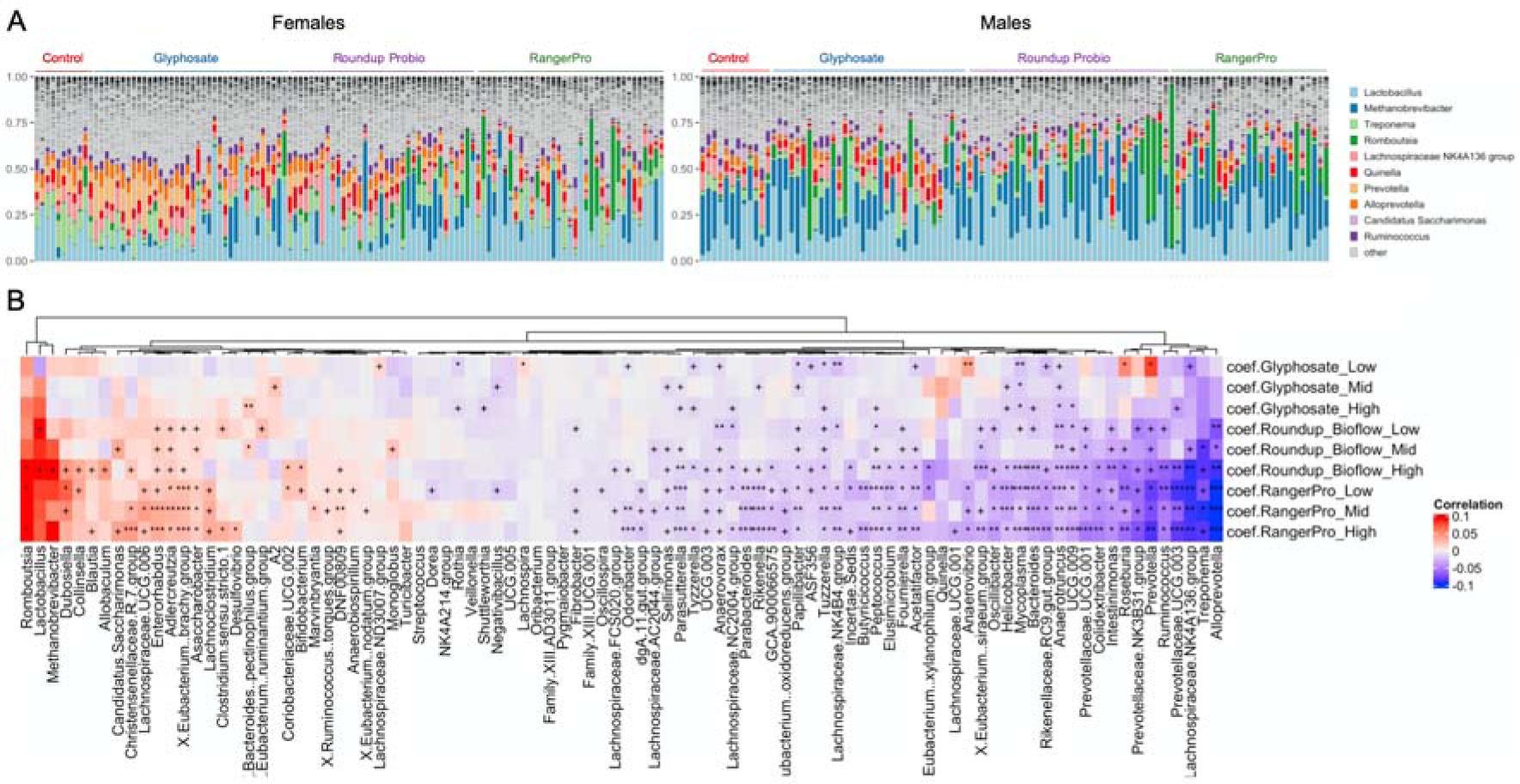
Glyphosate and its herbicide formulations Roundup Bioflow and RangerPro cause large scale alterations in gut microbiome taxonomic composition. Male and female Sprague-Dawley rats were exposed to glyphosate or the herbicide formulations at 0.5, 5 and 50mg/kg bw/day glyphosate starting at mid-gestation and ending at 13 weeks post-weaning. **A**. Visualisation of the top 10 genera across all the samples from the caecum microbiome dataset. **B**. Heat map of statistically significant differences in abundance caused by exposure to the test substances. The colour scale is the effect size from a multivariate association analysis showing if a bacterial genera is more abundant (red) or less abundant (blue) than in the control group. The statistical significance is the FDR indicated by symbols (*** FDR < 0.001 ; ** FDR < 0.01 ; * FDR < 0.05 ; FDR < 0.2).

Analysis of alterations in fungal composition at the genus level indicated that the abundance of *Kazachstania* (Ascomycota) was reduced while the abundance of *Gibberella, Penicillium, Claviceps, Cornuvesica, Candida, Trichoderma* and *Sarocladium* were increased by the treatments with the two Roundup formulated products (Figure 4). Glyphosate had limited effects on the mycobiome. Overall, these observations further suggest that exposure to glyphosate and its formulations affects the abundance of major bacteria taxa, which in turn reduces competition and allows opportunistic fungi to grow in the gut of the exposed animals.

**Figure 4.**
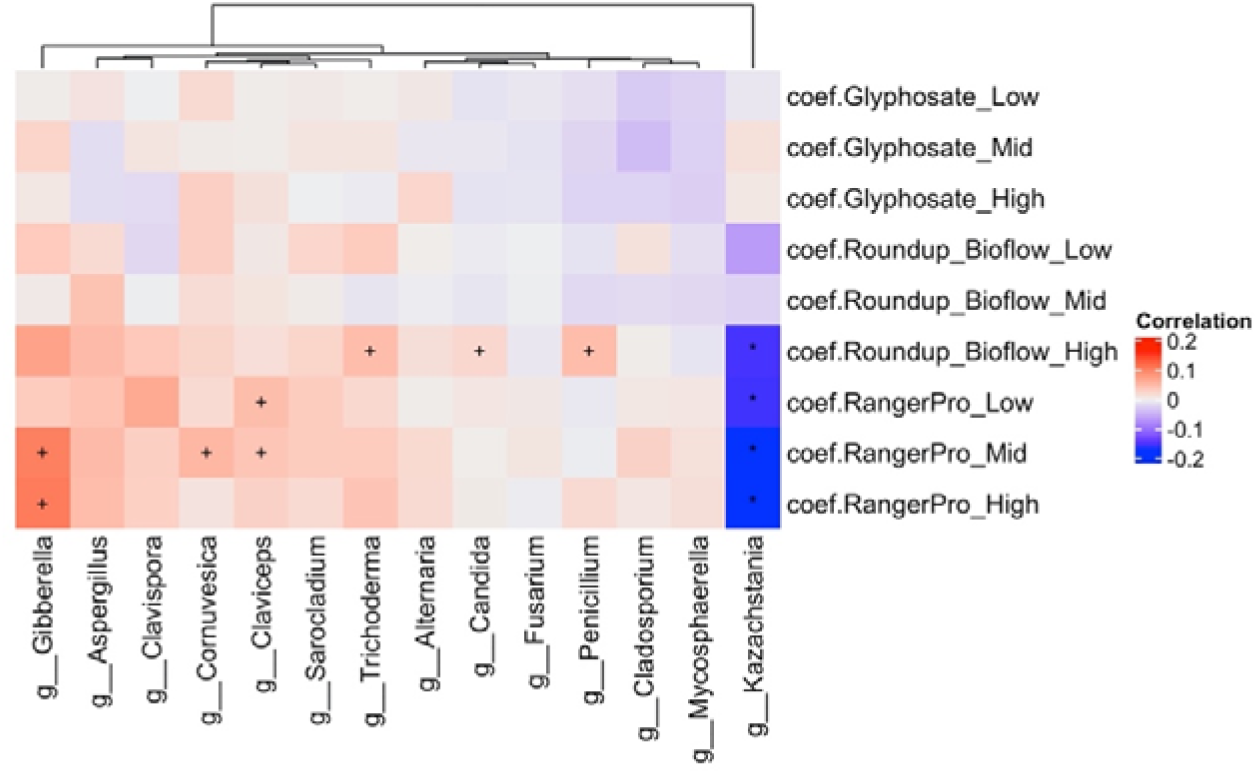
The formulations Roundup Bioflow and RangerPro alter the caecum mycobiome in rats. Male and female Sprague-Dawley rats were exposed to glyphosate or the herbicide formulations at 0.5, 5 and 50mg/kg bw/day glyphosate starting at mid-gestation and ending at 13 weeks post-weaning. The figure shows a heat map of statistically significant differences in abundance caused by the exposure to the tested herbicides in the caecum mycobiome. The colour scale is the effect size from a multivariate association analysis showing if a fungal genera is more (red) or less (blue) abundant than in the control group. The statistical significance is the FDR indicated by symbols (*** FDR < 0.001 ; ** FDR < 0.01 ; * FDR < 0.05 ; FDR < 0.2).

## Discussion

Here we have provided the most comprehensive description of the effects of glyphosate and two formulated glyphosate-based Roundup herbicide formulations on the bacterial and fungal composition of the caecum microbiome of rats. Our results provide information on the effects of glyphosate and two Roundup formulations on gut microbial composition at an unprecedented level of detail by combining an evaluation of fungal and bacterial composition on the same rat caecum samples.

Our previous study using a combination of shotgun metagenomics and metabolomics suggested that glyphosate does not have antibiotic properties, but in contrast could cause the proliferation of some bacteria that potentially use glyphosate as a source of phosphate (Mesnage et al. 2021c). The results from this study are very different, suggesting that glyphosate and two representative commercial Roundup formulations completely reshaped the rat gut microbiome. The current study started exposure doses prenatally which reveal effects of glyphosate that are not detected in adult animals with more mature and stable gut microbial communities. Starting treatment at a prenatal stage of development appears to be not only more representative of real-world exposure scenarios but is apparently able to reveal effects of glyphosate and Roundup formulations that are not detected when treatment is initiated in adult animals, which have more mature and stable gut microbial communities. The adult gut microbiome is relatively stable and resilient to environmental perturbations (Lozupone et al. 2012). In contrast, the developing gut microbiome in infants appears to be more sensitive to perturbations, which could durably impact health during adult life (Robertson et al. 2019).

Previous studies have suggested that *Lactobacilli*, and Firmicutes in general, are the bacteria most affected by glyphosate in the rat gut microbiome (Lozano et al. 2018; Mao et al. 2018; Nielsen et al. 2018). Although our results contradict these studies by showing increased abundance of different Firmicutes after glyphosate and Roundup exposure, it is interesting to note that a recent microevolutionary analysis of the EPSPS enzyme suggested that Firmicutes are significantly more resistant to glyphosate than other taxonomic groups (Rainio et al. 2021). It is possible that glyphosate exposure has favoured the growth of resistant strains of Firmicute species. However, they are not distinguishable with the present sequencing strategy and more focused metagenomic analyses will have to be performed to resolve this issue.

An important limitation in this study is that both 16S rRNA and ITS sequencing provided semiquantitative estimates of abundance. Each of the datasets was normalised to 100%, providing independent evaluations of bacterial and fungal abundance. However, fungi typically account for a small proportion of the gut microflora in comparison to bacteria representing approximately 0.1% of the total microbial load (Nash et al. 2017). In addition, not all of the single-celled prokaryotes are bacteria. Methanogenic archaea such as *Methanobrevibacter smithii* account for up to 10% of the human gut microbiome (Gaci et al. 2014). Small unicellular eukaryotes are also present. However, it is not clear if they are permanent residents of gut microbial ecosystems or if they do not colonise the gut durably. Our earlier study provided a global analysis of variations in the microorganism composition of the rat gut microbiome after treatment with glyphosate and Roundup Bioflow with shotgun metagenomics (Mesnage et al. 2021c). However, metagenomic applications in rats are also limited by the incompleteness of the taxonomic classification in gene catalogues and there is no gold standard to perform compositional analyses.

Few studies of the gut mycobiome have been conducted in rats. The dominant fungal taxa in our Sprague-Dawley colony were found to *Kazachstania pintolopesii*. This yeast was also found in laboratory mice received by the National Institutes of Health from different suppliers (Kurtzman et al. 2005). It is widespread and was even found to dominate the gut mycobiome of live-trapped mice from east Slovakia (Bendova et al. 2020). The species complex of Ascomycetous yeasts *Kazachstania* is found in rodents, humans, birds, horses, pigs, and cows, but *K. pintolopesii* is almost exclusively found in rodents (Kurtzman et al. 2005). Other fungi species were found at a limited abundance in our study. In another study, it was found that the gut microbiome of laboratory mice born to wild mice resembles that of wild mice with higher abundance of Ascomycota and a lower abundance of Basidiomycota compared to conventional laboratory mice (Rosshart et al. 2019). In humans, the mycobiome is low in diversity and mostly represented by yeast including *Saccharomyces, Malassezia*, and *Candida* (Nash et al. 2017). Altogether, our results on the rat gut mycobiome might be relevant for rodent populations but more studies in human samples should be done to extrapolate our findings to human populations. This can be performed using the SHIME® technology, which mimics the entire gastrointestinal tract by employing a series of reactors representing the different steps in food uptake and digestion (Venema and van den Abbeele 2013).

The changes in gut microbiome composition found in this rat study can have health consequences when they are found in human populations. Alpha diversity which is decreased by Roundup Bioflow and RangerPro is linked to human health, with lower levels of alpha diversity associated with chronic diseases(Manor et al. 2020). The health implications of the proliferation of various fungal species in the gut remains elusive, but it is worth noting that the fungal gut microbiome is mostly considered as a reservoir of pathogenic microbes which proliferate when the host is compromised leading to inflammation (Huffnagle and Noverr 2013). However, conclusions on health implications are limited by the methodology used in this study. It is important to note that health effects of bacteria are strain-dependent and bacteria from the same species can have very different effects. For instance, the probiotic *E. coli* Nissle 1917 strain can protect against the invasion by adherent-invasive *E. coli* B2 strains (Sassone-Corsi et al. 2016). It is thus not possible to definitely attribute beneficial or detrimental health effects on the basis of compositional changes at the genus levels.

The effects of glyphosate and its formulated Roundup products in our study were in general comparable although patterns of statistical significance were variable. The sex of the host is known to shape its gut microbiota via the effects of sex hormones both in rodents and in humans (Valeri and Endres 2021). Sex-dependent gut microbiota differences in response to dietary interventions and exposures to toxic chemicals are known in diverse animal models and humans (Valeri and Endres 2021). Sex-specific effects of glyphosate formulations on gut microbiota was suggested in another study but the limited number of samples available in this investigation prevented drawing any general conclusions (Lozano et al. 2018).

Since applicators always spray a mixture of glyphosate and co-formulants, it is relevant to assess the toxicity of the complete formulated herbicides (not glyphosate alone) to understand health effects of their application. Our results show that the formulated products Roundup Bioflow and RangerPro caused more alterations than glyphosate alone at the same glyphosate-equivalent doses. It is likely that the surfactants included in Roundup Bioflow and RangerPro contribute to the gut microbial alterations. However, the changes caused by glyphosate and the two formulated products are comparable but more severe with the formulations, suggesting that the surfactants may enhance the effects of glyphosate as they do in plants in order for glyphosate to act as an efficient weedkiller. Numerous studies have shown that POEA surfactants contribute to the toxicity of glyphosate weedkillers. Studies showing that POE 15 tallow amine was more toxic than glyphosate have been available since the end of the 1970’s (Folmar et al. 1979). The formulation MON 2139 containing POE 15 tallow amine was 10 to 40 times more toxic than glyphosate in different fish species (Folmar et al. 1979; Wan et al. 1989). We have previously shown that Roundup Bioflow, which does not contain POEA but a quaternary ammonium surfactant (Mesnage et al. 2021b) causes more gut microbiome alterations than glyphosate alone in rats (Mesnage et al. 2021c). Although studies have described the presence of POEA surfactants in RangerPro (Mesnage et al. 2021a), the complete co-formulant profile remains unknown. Further studies testing the surfactants alone will be required to understand if they can be a source of effects in the absence of glyphosate.

In conclusion, we reveal that early life exposures starting prenatally to glyphosate or its formulated products Roundup Bioflow and RangerPro cause large changes in the composition of the rat gut microbiota. This indicates that alterations in gut microbiome composition will have to be taken into account in the next phases of the *Global Glyphosate Study* addressing long-term toxicity, carcinogenicity and multi-generational effects of glyphosate and glyphosate -based herbicide formulations.

## Acknowledgements

The gut microbiome analysis was funded by the Sustainable Food Alliance (USA), whose support is gratefully acknowledged. The Global Glyphosate Study was funded by the Ramazzini Institute (Italy), the Heartland Health Research Alliance (USA), the Boston College (USA), the Fondazione Carisbo (Italy), the Fondazione del Monte di Bologna e Ravenna (Italy), the Coop Reno (Italy) and the Coopfond Fondo Mutualistico Legacoop (Italy).

## References

Bailey DC, Todt CE, Burchfield SL, et al. (2018) Chronic exposure to a glyphosate-containing pesticide leads to mitochondrial dysfunction and increased reactive oxygen species production in Caenorhabditis elegans. Environ Toxicol Pharmacol 57:46–52

Benbrook CM (2016) Trends in glyphosate herbicide use in the United States and globally. Environmental Sciences Europe 28(1):3

Benbrook CM (2019) How did the US EPA and IARC reach diametrically opposed conclusions on the genotoxicity of glyphosate-based herbicides? Environmental Sciences Europe 31(1):2

Bendova B, Pialek J, Dureje L, et al. (2020) How being synanthropic affects the gut bacteriome and mycobiome: comparison of two mouse species with contrasting ecologies. BMC Microbiol 20(1):194

Boocock MR, Coggins JR (1983) Kinetics of 5-enolpyruvylshikimate-3-phosphate synthase inhibition by glyphosate. FEBS Lett 154(1):127–33

Duforestel M, Nadaradjane A, Bougras-Cartron G, et al. (2019) Glyphosate Primes Mammary Cells for Tumorigenesis by Reprogramming the Epigenome in a TET3-Dependent Manner. Front Genet 10:885

Folmar LC, Sanders HO, Julin AM (1979) Toxicity of the herbicide glyphosphate and several of its formulations to fish and aquatic invertebrates. Arch Environ Contam Toxicol 8(3):269–78

Gaci N, Borrel G, Tottey W, O’Toole PW, Brugère J-F (2014) Archaea and the human gut: new beginning of an old story. World journal of gastroenterology 20(43):16062–16078

Gomes MP, Juneau P (2016) Oxidative stress in duckweed (Lemna minor L.) induced by glyphosate: Is the mitochondrial electron transport chain a target of this herbicide? Environ Pollut 218:402–409 doi:10.1016/j.envpol.2016.07.019

Heap I (2014) Global perspective of herbicide-resistant weeds. Pest Manag Sci 70(9):1306–15

Huffnagle GB, Noverr MC (2013) The emerging world of the fungal microbiome. Trends in microbiology 21(7):334–341

Huseyin CE, O’Toole PW, Cotter PD, Scanlan PD (2017) Forgotten fungi—the gut mycobiome in human health and disease. FEMS Microbiol Rev 41(4):479–511

Kurtzman CP, Robnett CJ, Ward JM, Brayton C, Gorelick P, Walsh TJ (2005) Multigene phylogenetic analysis of pathogenic candida species in the Kazachstania (Arxiozyma) telluris complex and description of their ascosporic states as Kazachstania bovina sp. nov., K. heterogenica sp. nov., K. pintolopesii sp. nov., and K. slooffiae sp. nov. J Clin Microbiol 43(1):101–11

Lozano VL, Defarge N, Rocque LM, et al. (2018) Sex-dependent impact of Roundup on the rat gut microbiome. Toxicol Rep 5:96–107

Lozupone CA, Stombaugh JI, Gordon JI, Jansson JK, Knight R (2012) Diversity, stability and resilience of the human gut microbiota. Nature 489(7415):220–230

Maggi F, Tang FHM, la Cecilia D, McBratney A (2019) PEST-CHEMGRIDS, global gridded maps of the top 20 crop-specific pesticide application rates from 2015 to 2025. Scientific Data 6(1):170

Manor O, Dai CL, Kornilov SA, et al. (2020) Health and disease markers correlate with gut microbiome composition across thousands of people. Nature Communications 11(1):5206

Mao Q, Manservisi F, Panzacchi S, et al. (2018) The Ramazzini Institute 13-week pilot study on glyphosate and Roundup administered at human-equivalent dose to Sprague Dawley rats: effects on the microbiome. Environmental Health 17:50

McMurdie PJ, Holmes S (2013) phyloseq: an R package for reproducible interactive analysis and graphics of microbiome census data. PLoS One 8(4):e61217

Mesnage R, Antoniou MN (2020) Computational modelling provides insight into the effects of glyphosate on the shikimate pathway in the human gut microbiome. Current Research in Toxicology 1:25–33

Mesnage R, Benbrook C, Antoniou MN (2019) Insight into the confusion over surfactant co-formulants in glyphosate-based herbicides. Food Chem Toxicol 128:137–145

Mesnage R, Defarge N, Spiroux de Vendomois J, Seralini GE (2015) Potential toxic effects of glyphosate and its commercial formulations below regulatory limits. Food Chem Toxicol 84:133–53

Mesnage R, Ferguson S, Mazzacuva F, Caldwell A, Halket J, Antoniou MN (2021a) Cytotoxicity mechanisms and composition of the glyphosate formulated herbicide RangerPro. BIORXIV 469091.

Mesnage R, Mazzacuva F, Caldwell A, Halket J, Antoniou MN (2021b) Urinary excretion of herbicide co-formulants after oral exposure to roundup MON 52276 in rats. Environmental Research 197:111103

Mesnage R, Phedonos A, Biserni M, et al. (2017) Evaluation of estrogen receptor alpha activation by glyphosate-based herbicide constituents. Food Chem Toxicol 108(Pt A):30–42

Mesnage R, Teixeira M, Mandrioli D, et al. (2021c) Use of shotgun metagenomics and metabolomics to evaluate the impact of glyphosate or Roundup MON 52276 on the gut microbiota and serum metabolome of Sprague-Dawley rats.. Environmental Health Perspectives Jan;129(1):17005.

Nash AK, Auchtung TA, Wong MC, et al. (2017) The gut mycobiome of the Human Microbiome Project healthy cohort. Microbiome 5(1):153 doi:10.1186/s40168-017-0373-4

Nielsen LN, Roager HM, Casas ME, et al. (2018) Glyphosate has limited short-term effects on commensal bacterial community composition in the gut environment due to sufficient aromatic amino acid levels. Environmental Pollution 233:364–376

Olorunsogo OO (1990) Modification of the transport of protons and Ca2+ ions across mitochondrial coupling membrane by N-(phosphonomethyl)glycine. Toxicology 61(2):205–9

Pereira AG, Jaramillo ML, Remor AP, et al. (2018) Low-concentration exposure to glyphosate-based herbicide modulates the complexes of the mitochondrial respiratory chain and induces mitochondrial hyperpolarization in the Danio rerio brain. Chemosphere 209:353–362

Portier CJ (2020) A comprehensive analysis of the animal carcinogenicity data for glyphosate from chronic exposure rodent carcinogenicity studies. Environmental Health 19(1):18

Rainio MJ, Ruuskanen S, Helander M, Saikkonen K, Saloniemi I, Puigbò P (2021) Adaptation of bacteria to glyphosate: a microevolutionary perspective of the enzyme 5-enolpyruvylshikimate-3-phosphate synthase. Environmental Microbiology Reports 13(3):309–316

Robertson RC, Manges AR, Finlay BB, Prendergast AJ (2019) The Human Microbiome and Child Growth - First 1000 Days and Beyond. Trends Microbiol 27(2):131–147

Robinson C, Portier CJ, ČAvoŠKi A, et al. (2020) Achieving a High Level of Protection from Pesticides in Europe: Problems with the Current Risk Assessment Procedure and Solutions. European Journal of Risk Regulation:1–31

Rosshart SP, Herz J, Vassallo BG, et al. (2019) Laboratory mice born to wild mice have natural microbiota and model human immune responses. Science 365(6452)

Sassone-Corsi M, Nuccio S-P, Liu H, et al. (2016) Microcins mediate competition among Enterobacteriaceae in the inflamed gut. Nature 540(7632):280

Tsiaoussis J, Antoniou MN, Koliarakis I, et al. (2019) Effects of single and combined toxic exposures on the gut microbiome: Current knowledge and future directions. Toxicol Lett 312:72–97

Valeri F, Endres K (2021) How biological sex of the host shapes its gut microbiota. Front Neuroendocrinol 61:100912

Venema K, van den Abbeele P (2013) Experimental models of the gut microbiome. Best Pract Res Clin Gastroenterol 27(1):115–26

Wan MT, Watts RG, Moul DJ (1989) Effects of different dilution water types on the acute toxicity to juvenile Pacific salmonids and rainbow trout of glyphosate and its formulated products. Bull Environ Contam Toxicol 43(3):378–85

Wise T, MacDonald GJ, Klindt J, Ford JJ (1992) Characterization of thymic weight and thymic peptide thymosin-beta 4: effects of hypophysectomy, sex, and neonatal sexual differentiation. Thymus 19(4):235–44

Woźniak E, Reszka E, Jabłońska E, Balcerczyk A, Broncel M, Bukowska B (2020) Glyphosate affects methylation in the promoter regions of selected tumor suppressors as well as expression of major cell cycle and apoptosis drivers in PBMCs (in vitro study). Toxicology in Vitro 63:104736

